# A combinatorial biomolecular strategy to identify peptides for improved transport across the sputum of cystic fibrosis patients and the underlying epithelia

**DOI:** 10.1101/659540

**Authors:** Jasmim Leal, Xinquan Liu, Xiujuan Peng, Rashmi P. Mohanty, Dhivya Arasappan, Dennis Wylie, Sarah H. Schwartz, Jason J. Fullmer, Bennie C. McWilliams, Hugh D. C. Smyth, Debadyuti Ghosh

**Author notes:** Corresponding author Debadyuti Ghosh.

## Abstract

Drugs and drug delivery systems have to traverse multiple biological barriers to achieve therapeutic efficacy. In diseases of mucosal-associated tissues such as cystic fibrosis (CF), successful delivery of gene and drug therapies remains a significant challenge due to an abnormally concentrated viscoelastic mucus, which prevents ~99% of all drugs and particles from penetrating the mucus barrier and the underlying epithelia for effective therapy, resulting in decreased survival. We used combinatorial peptide-presenting phage libraries and next-generation sequencing to identify hydrophilic, close to net-neutral charged peptides that penetrate the mucus barrier *ex vivo* in sputum from CF patients with ~600-fold better penetration than a positively charged control. After mucus penetration, nanoparticles conjugated with our selected peptides successfully translocated into lung epithelial cells derived from CF patients and demonstrated up to three-fold improved cell uptake compared to non-modified carboxylated- and gold standard PEGylated-nanoparticles. The selected peptides act as surface chemistries with synergistic functions to significantly improve the ability of drug delivery systems to overcome the human mucosal barriers and provide efficient cellular internalization. Our screening strategy provides a biologically-based discovery assay that directly addresses transport through mucus and cell barriers and has the potential to advance drug and gene delivery to multiple mucosal barriers.

## Introduction

For successful treatment of mucosal-associated diseases, such as cystic fibrosis (CF), HIV, asthma, and chronic obstructive pulmonary disease, it is critical for drug and gene therapies to cross several biological barriers to enter the target cells at therapeutic concentrations. One of the initial and primary barriers is the protective mucus layer lining the epithelia in the eyes, airways, gastrointestinal and cervicovaginal tracts [1–3]. Mucus is a complex biopolymer composed mainly of water, mucin glycoproteins, lipids, DNA, non-mucin proteins, salts, and cell debris [4]. The mucus layer acts as a transport barrier and can greatly hinder the diffusion of drugs, particles, and other molecules [5–7] via size filtering, mucociliary clearance, and intermolecular interactions, including electrostatic, hydrophobic, and/or other specific binding interactions [8]. For example in CF, the hyperconcentrated mucus layer traps foreign pathogens, resulting in chronic bacterial infections and concomitantly prevents up to 99% of drugs and particles from penetration needed for successful therapy [9, 10]. As a result, it is a long-standing challenge for drugs to permeate the mucus barrier and reach the underlying epithelia.

Considerable advancements have been made towards the development of different drug delivery strategies that attempt to overcome the mucus barrier, including use of mucolytic agents [11, 12], mucoadhesive [13–15], and mucus-penetrating [16–19] delivery systems. It has been demonstrated that the use of net-neutral charge hydrophilic polymers can improve the transport of nanoparticles and minimize interactions with respiratory mucus [17, 20], gastrointestinal mucus [21, 22], and cervical mucus [23–25]. However, it has been estimated that only 35% of 200 nm nanoparticles functionalized with poly(ethylene glycol) (PEG) 3.4 kDa are capable of penetrating through a 10 μm CF sputum layer within 20 min [17]. Also, there have been numerous clinical studies that have demonstrated that people have pre-existing anti-PEG antibodies [26, 27] and present an immune response after the administration of PEGylated proteins [26, 28], aptamers [29, 30], and nanoparticles [31, 32]. As a result, there are potential concerns for the safety and feasibility of PEG carriers prior to their use for mucus penetrating delivery, and there is a need to find alternative mucus-penetrating chemistries.

After mucus penetration, drug delivery systems must enter target cells to efficiently deliver their therapeutic payload. Although PEGylated systems demonstrate improved diffusion in mucus via decreased hydrophobic and electrostatic interactions, PEGylation can hinder cellular uptake of the drug carrier system [33–37]. PEGylation has been shown to increase serum protein binding of lipid/DNA complexes *in vitro* and decrease gene delivery *in vivo* [38, 39]. In genetic diseases such as CF, it is essential that delivery systems can deliver small molecules or nucleic acid therapeutics intracellularly. For enhanced cellular uptake of carriers, targeting ligands such as peptides have been tested. For example, poly(lactic-co-glycolic acid) PLGA nanoparticles modified with cell-penetrating peptides through a PEGylated phospholipid linker have been tested for gene delivery to the lungs, with modest gene correction effects (<1%) [40]. Also, PEGylated cell-penetrating peptide nanoparticles have been tested *in vitro* and *in vivo* for lung gene therapy with different densities of PEG coatings, however a decrease in DNA uptake and transgene expression was found with increasing PEGylation rates, compared to non-PEGylated particles [41]. Current strategies mostly focus on either mucus penetration or cell penetration, but there has not been a single surface chemistry that addresses both mucus and cellular barriers.

However, in nature, viruses have evolved to possess properties for favorable interactions for mucus transport [42, 43]. Complex virus-like particles that are hydrophilic and display highly densely charged, alternating cationic and anionic amino acids on their surface have demonstrated unhindered, diffusive transport through mucus comparable to saline [42]. It has also been demonstrated that peptides with asymmetric charge properties had improved transport in reconstituted mucin [44]. Collectively, these results suggest that biomolecules have diverse physicochemical properties (i.e. charge, hydrophilicity, their spatial distribution) and by changing their permutation, a large molecular diversity can be achieved and leveraged to identify new surface coatings with favorable interactions for mucus-penetrating transport.

In this work, we identify peptide coatings with the ability to penetrate CF mucus using phage-assisted directed evolution and ultimately, achieve intracellular uptake in mutant CF epithelial cells. Bacteriophage (phage), or viruses that infect bacteria, have been engineered as display systems to express combinatorial libraries of up to 10^9^ different peptides or proteins on their viral coat proteins (i.e. different peptide per phage), which function as a collection of surface chemistries with diverse physicochemical properties and functionalities [45]. Iteratively screening of peptide-presenting phage libraries is a powerful high-throughput method for identification of peptide coatings to a variety of targets with no known lead structures or substrates that can favorably interact with mucus for penetration and desired functionalities. The highly diverse libraries are collapsed to a few leads by performing repetitive rounds of selection against the target, or selection pressure (i.e. biopanning) [46, 47].

Here, we screened peptide-presenting phage libraries against a mucus barrier, used next-generation DNA sequencing to identify the genetically-encoded peptides from a vast sequence space, and then validated their mucus-penetration *ex vivo* using clinical samples from the sputum of cystic fibrosis patients. In CF, the mutation of the CF transmembrane conductance regulator gene expressed in epithelial cells [48, 49] manifests in unbalanced ion transport, impaired mucociliary clearance, dehydration of the airway epithelia, and the accumulation of abnormal, hyperconcentrated mucus [50]. The increase in mucus concentration and inability to clear the mucus creates a physical barrier that traps pathogens and drastically decreases drug diffusion, absorption, bioavailability, and consequently limiting therapeutic outcomes of mucosal drug delivery systems [51, 52]. In this study, we present a general strategy to identify mucus-penetrating peptides, validate identified peptides in CF sputum samples from patients, and confirm the cellular uptake of nanoparticles functionalized with mucus-penetrating peptides. This biomolecule-based approach can be translated to therapeutic delivery strategies of drug and gene carriers across mucosal barriers and the dense mucus barrier present in disease states like cystic fibrosis, chronic pulmonary obstructive disease, and asthma.

## Results

### High-throughput sequencing of phage libraries screened against CF mucus indicates a convergence for unique peptides

The constructed T7 phage CX7C library had a diversity (i.e. number of phage displaying unique peptide sequences) of 3.7×10^5^ distinct clones, as determined by next-generation sequencing (Supplementary Table 2) and a phage concentration of 1.24 x 10^11^ plaque forming units per mL (pfu/mL), as determined by standard double-layer plaque assay. To identify peptide-presenting phage that diffuse across the barrier, the T7 phage CX7C library was iteratively screened against a CF mucus model (Fig. 1A). One hundred microliters of CF mucus were added on the donor side of the transwell system, which contained 600 μL of 1X phosphate buffered saline (PBS) on the receiver compartment. In the first round, 4.2 × 10^9^ pfu of the original phage library was added on top of the CF mucus layer, and the number of phage that diffused to the receiver side after 1 hour was quantified by plaque assay. Ten microliter aliquots were also collected after 15 and 30 minutes for further analysis by high-throughput sequencing. After 1 hour, the entire eluate was collected, amplified (i.e. grow more copies), and was incubated on CF mucus for a second round of screening. This procedure was repeated for a total of three rounds of screening.

**Figure 1.**
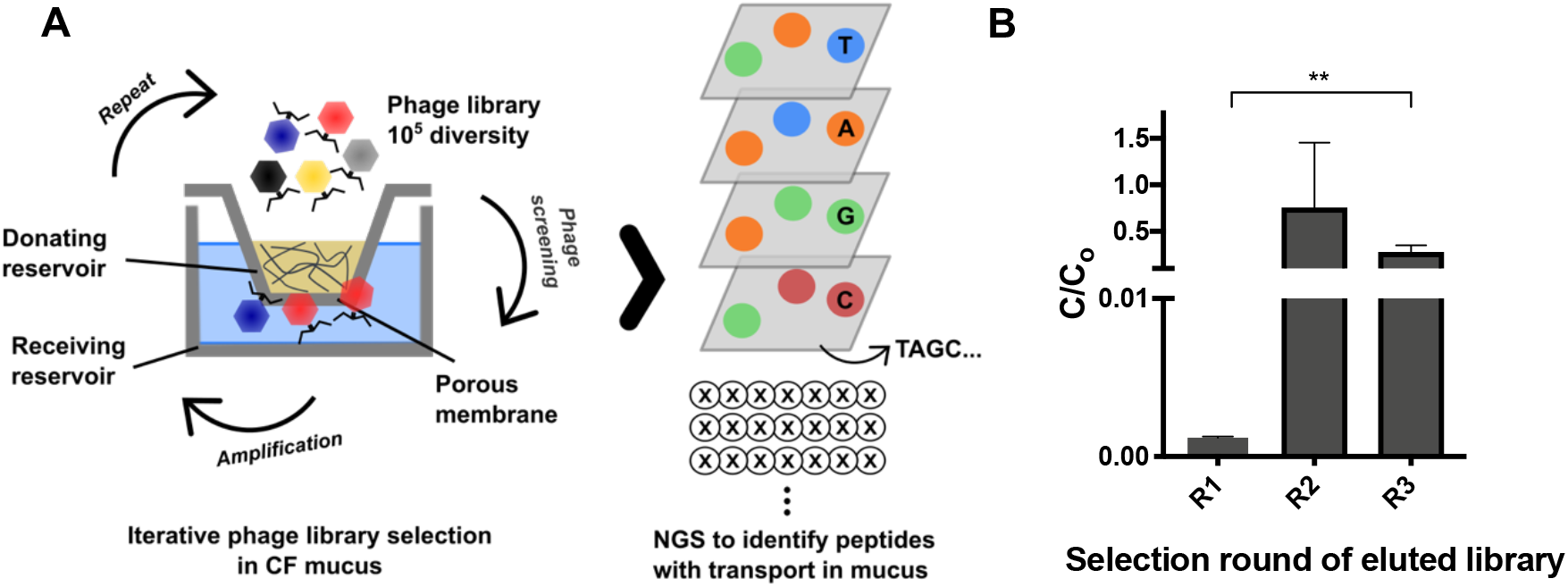
Overview of the iterative phage library selection in CF mucus model. (A) A T7 phage peptide library with an initial diversity of 10^5^ distinct clones is added on top of a CF mucus layer in a Transwell system (Corning, MA) and incubated in the donating reservoir with 1X PBS. Phage that penetrate through CF mucus are collected, quantified by standard double-layer plaque assay, and amplified by E. coli for use either in the next selection round or subjected to PCR and next-generation high-throughput DNA sequencing. (B) Enrichment of phage plaque forming units recovered from the basolateral side over three rounds of screening (R1 to R3, n = 3), quantified by standard double-layer plaque assay (unpaired, twotailed Student’s t-test, p<0.01).

There was a significant 232-fold increase in the output to input of phage (C/C0) that penetrated through CF mucus from the third round compared to the first round (Fig. 1B). This increase under selection pressure of mucus barrier suggests there is possible enrichment for clones with desired properties (i.e. mucus penetration) in CF mucus [47, 53].

Next, the phage DNA from all three selection rounds in replicates was isolated, PCR amplified, and sequenced by high-throughput sequencing. For each of the samples, an average of 635,004 ± 124,391 DNA sequences was obtained. From these sequences, sequences that did not encode for the CX7C peptide sequence or included stop codons before the first cysteine residue were excluded from subsequent analysis and experiments; otherwise, reads would contain a significant fraction of sequences with TAG stop codons or not display any peptide sequence. The number of unique peptide sequences in each sample decreased considerably with each subsequent round of selection (Supplementary Table 2), which has been observed by others [53, 54].

To determine the distribution and abundance of peptide sequences and reproducibility (i.e. similarity between replicates) with each round of selection, we analyzed density plots of peptide frequencies (i.e. counts) (Fig. S1). With each successive round of selection (denoted as R1-R3), the frequencies of specific peptide sequences in both replicates (denoted as replicate 1 and replicate 2) were reproducible; peptides observed with 100 counts in replicate 1 were observed at similar frequencies in replicate 2. Moreover, from the heat map, there is a convergence of the most frequent peptides. With each successive round of selection, the number of unique sequences decreases, and there is a convergence of abundant peptides (Fig. S1), which suggests there is selection and enrichment for unique peptide sequences [47].

Next, the amino acid composition of peptide sequences from each round of selection was compared to the original library (Fig. S2). Here, mostly hydrophobic amino acids had an overall decrease in their frequency during selection compared to the original library. There was an enrichment in glycine (G), serine (S), and in the acidic residues glutamic acid (E) and aspartic acid (D) over rounds compared to the original library. Basic residues such as histidine (H), lysine (K), and arginine (R) had an overall decrease in frequency after three rounds of selection against CF mucus.

### Identification of CF mucus-penetrating peptides by high throughput sequencing leads to negative to neutral net-charge and hydrophilic sequences

After high-throughput sequencing of DNA isolated from phage eluates, sequences were translated, and the top 20 most abundant peptide sequences and their physicochemical properties (i.e. charge and hydropathy GRAVY score) were calculated (Fig. 2). The median net-charge was close to −1.0 in the three rounds; however, the interquartile range indicates there is a trend for the distribution to approach neutral charge with each round of selection. The GRAVY score gradually increased with subsequent rounds of selection, but overall, sequences were mostly hydrophilic to neutral.

**Figure 2.**
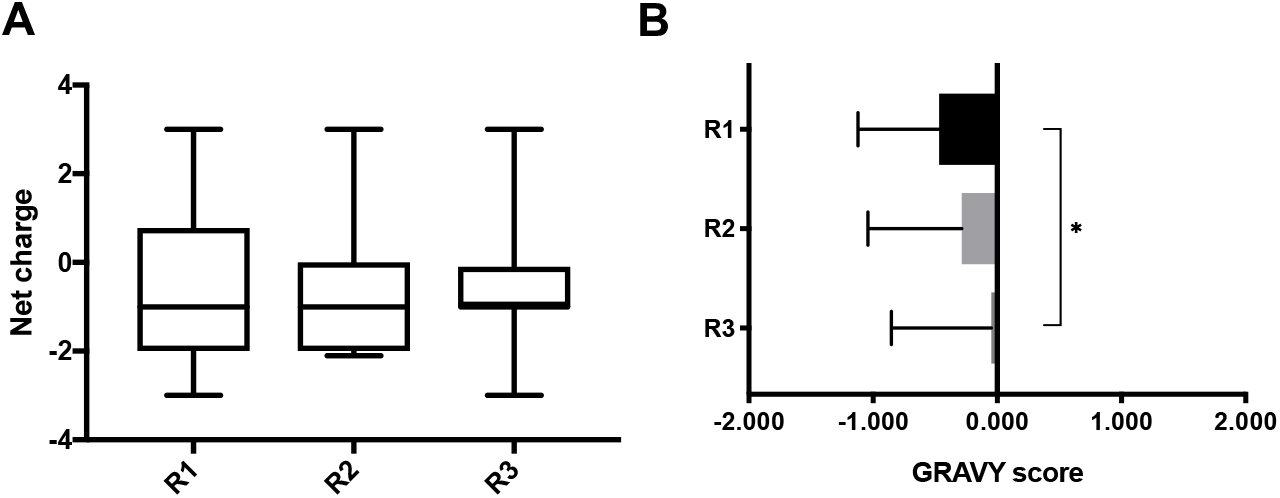
(A) Net charge at pH 7.0 and (B) GRAVY score of top 20 peptide hits in 60 minutes eluates, found by high-throughput sequencing of phage display selection against CF mucus. Net charge was calculated by Protein Calculator v3.4 (The Scripps Research Institute, La Jolla, CA. Available at http://protcalc.sourceforge.net/), and GRAVY score was calculated according to the literature [55]. (A) median, IQR. (B) mean ± SD. One-way analysis of variance (ANOVA) with Tukey’s multiple comparisons test (p<0.05).

Next, BLASTp local alignment search was done on the top 20 most abundant peptide sequences to identify their sequence to an existing database of known proteins. Of interest, peptide sequences PEPTIDE-2, PEPTIDE-3, and PEPTIDE-4 had homology to mucin 16 (MUC16), mucin 2 precursor (MUC2), and mucin 19 precursor (MUC19), respectively (Table 1). MUC16 is a cell surface associated mucin present in the airways, salivary glands, cervix, and eyes; MUC2 and MUC19 are secreted, gel-forming mucins present either in the airways, salivary glands, intestine, cervix, and/or eyes [2]. Another sequence of interest, PEPTIDE-1, was present in the top 20 most abundant peptide sequences throughout all rounds and in sequenced replicates. The aforementioned sequences possess either slightly negative or close to neutral net charges at pH 7.0 and were close to neutral or hydrophilic as indicated by the calculated GRAVY index score (Table 1) [55].

**Table 1.**
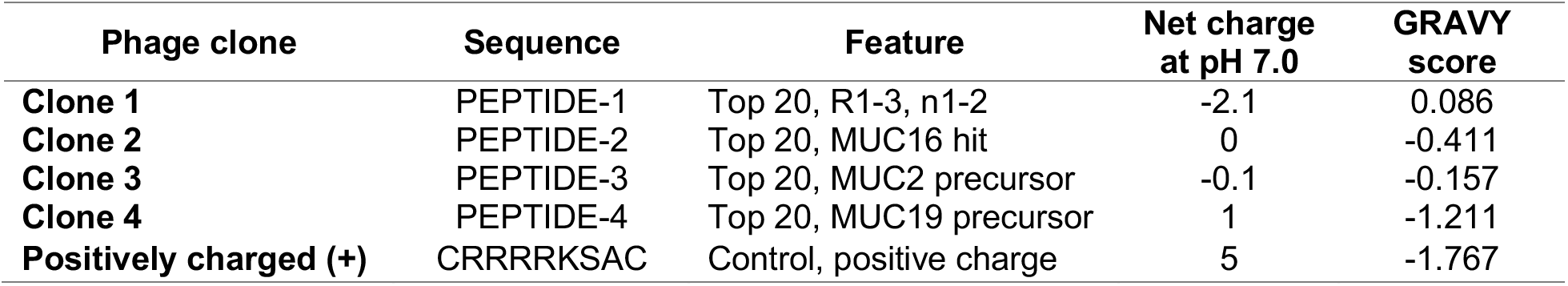
Physicochemical properties of selected mucus-penetrating peptides. Amino acid sequences, net charge at 7.0, and GRAVY score of peptide hits found by high-throughput sequencing of phage display selection against CF mucus. Net charge was calculated by Protein Calculator v3.4 (The Scripps Research Institute, La Jolla, CA. Available at http://protcalc.sourceforge.net/), and GRAVY score was calculated according to the literature [55].

To ensure that the frequency of identified sequences was due to screening and not growth bias, these sequences were compared to the naïve library grown without selection [53, 56]. The naïve phage library was amplified without any selection pressure for three rounds, and these samples were sequenced by high-throughput sequencing to identify overrepresented sequences indicative of fast growers. Selected peptide sequences from Table 1 were not amongst the overrepresented sequences present in the amplified rounds of the naïve library. In addition, we compared the selected peptide sequences to databases that have identified sequences with growth bias and/or non-specific affinity, also known as “target-unrelated peptides” (TUP) (54, 55); the selected peptide sequences (Table 1) were not found in these comprehensive databases. These results indicate that selected peptides are not artefactual and can be further validated for penetrating transport through the mucus barrier.

### Permeation assays of phage clones displaying the identified peptides indicates selection for phage particles with enhanced diffusion across CF mucus

#### Permeation assay across CF mucus model

We performed a permeation assay in a transwell system similar to the phage screening to confirm that identified phage-displayed peptides (Table 1) improve diffusion of phage across the CF mucus model. Each phage clone (2.0 × 10^9^ genomic copies (gc)) displaying PEPTIDE-1, PEPTIDE-2, PEPTIDE-3, or PEPTIDE-4 (denoted as Clones 1-4, respectively), along with a positive charge phage control (Clone (+)) was added on top of a CF mucus layer on the donor compartment of a transwell system. Phage that diffused across the mucus layer were recovered from the receiver compartment, and the amount of phage (Q) in the eluates was quantified by qPCR. As shown in Fig. 3A, phage clones 3 and 4 permeated across CF mucus 9- and 13-fold higher, respectively, compared to the positively charged control clone (+). Clone 1 did not show improved permeation compared to control, and clone 2 showed modest 3-fold improved permeation compared to the positively charged control clone (+). These findings suggest that the identified clones 3 and 4 have the ability to facilitate phage diffusion across CF mucus.

**Figure 3.**
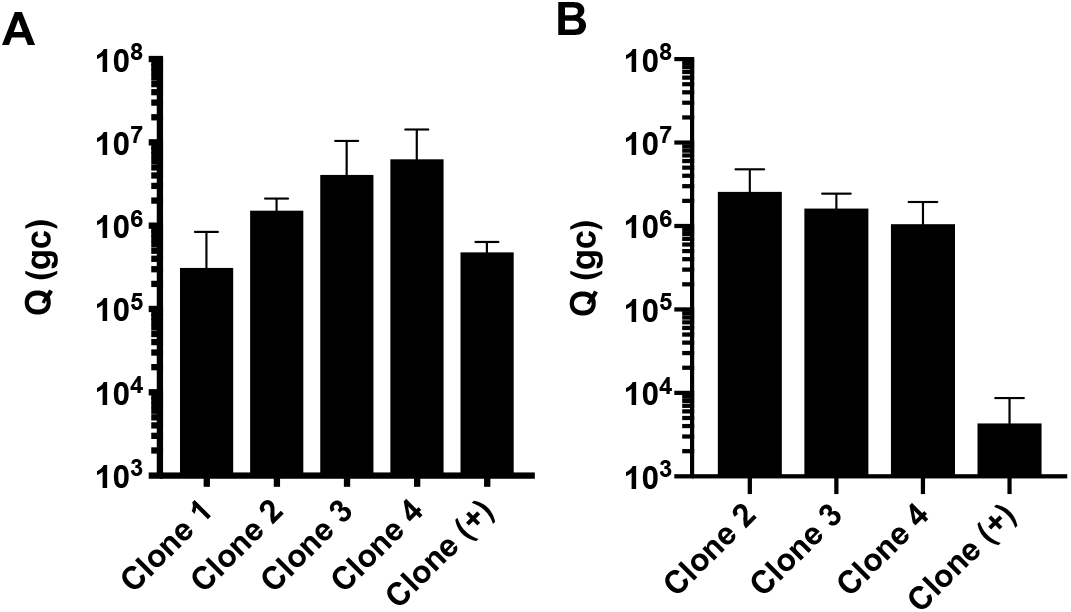
Amount of phage clones that permeate across (A) CF mucus model, and (B) CF sputum. Each phage clone (2.0 × 10^9^ genomic copies (gc)) was added on top of a CF mucus layer on the donor compartment of a transwell system. Phage that diffused across the mucus layer were recovered from the basolateral side at 2 hours, and phage amounts (Q) in the eluates were determined by qPCR (ns, not significant, p=0.1000 for clone 4 and clone (+) in CF mucus; p=0.1371 for clone 2 and clone (+) in CF sputum). Data represents mean ± SD (n = 3). Kruskal-Wallis test with Dunn’s multiple comparisons was used.

#### Permeation assay of phage-displayed peptides against pooled CF sputum from patients

Based on our permeation studies of phage-displayed peptides against CF-like mucus, we selected the best phage-displayed peptides and validated their ability to improve diffusion of phage across patient samples. Here we performed a permeation assay in a transwell system similar to the CF-like mucus permeation assay but used CF sputum from patients (n = 15) pooled together to minimize patient-to-patient variation. Phage were incubated and quantified similar to the CF-like permeation assay. As shown in Fig. 3B, phage clones 2, 3, and 4 permeated across CF sputum 590-, 375-, and 244-fold more than the positively charged control clone (+), respectively.

#### Permeation assay across CF sputum from individual patients

To determine patient-to-patient differences in diffusion of phage clones validated in CF mucus, we performed a permeation assay against CF sputum from individual patients (N=6). Fig. 4 shows the amount of each clone that permeates through the sputum of individual patients. There is considerable variability in phage permeability amongst patients, regardless of phage clone used. In individual patient samples, phage that permeated was independent of the peptide sequence—all tested phage clones either diffused through the barrier or demonstrated poor permeability. This suggests that phage diffusion in CF sputum might be patient-specific and can be governed by different properties in sputum, such as composition, solids content, and pore size.

**Figure 4.**
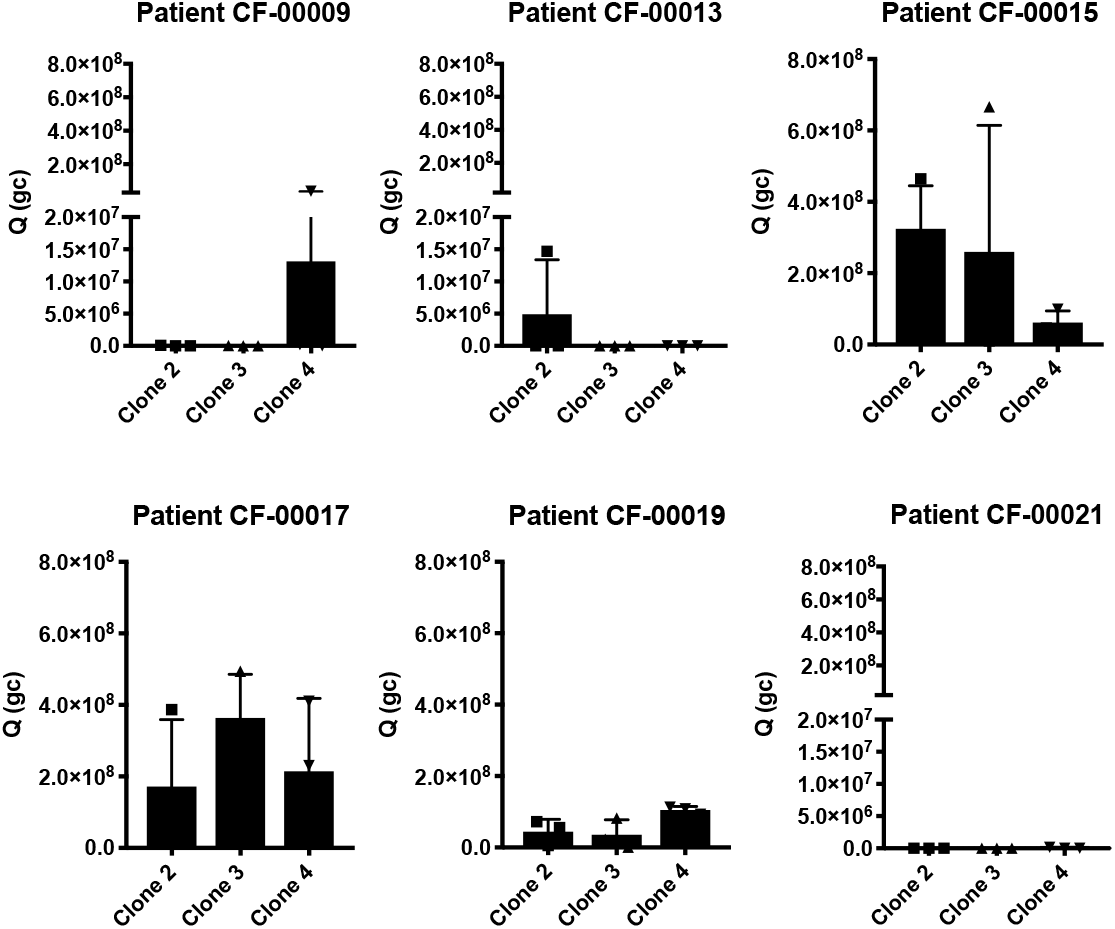
Phage clones permeation across CF sputum from individual patients. Each phage clone (2.0 × 10^9^ genomic copies (gc)) was added on top of a CF sputum layer on the donor compartment of a transwell system. Phage that diffused across the CF sputum were recovered from the basolateral side at 2 hours, and phage amounts (Q) in the eluates were determined by qPCR. Data represents mean ± SD (n = 3), Kruskal-Wallis test with Dunn’s multiple comparisons, not statistically significant.

### Biochemical characterization of CF sputum indicates patient heterogeneity

To characterize the components of CF sputum that could affect phage permeation, we measured the solids content, disulfide bond (i.e. cystine), mucin, and DNA concentration, and we quantified the porosity (i.e. void area %) of CF sputum samples. As shown in Fig. 5A, sputum samples from patients CF-00017, CF-00019, and CF-00021 had from 2- to 7-fold statistically significant higher solids content compared to the other patient samples (p<0.0001). The cystine content of the CF pooled sample was approximately 2- to 3-fold lower compared to individual patients, and patients CF-00017, CF-00019, and CF-00021 had the highest cystine content among all the samples tested (Fig. 5B). In Fig. 5C, patients CF-0009 and CF-00015 had lowest DNA concentrations, with approximately 60-fold lower concentrations compared to patients CF-00017, CF-00019, and CF-00021. Mucin concentration was 2-fold higher in patients CF-00017, CF-00019, and CF-00021, compared to the other samples, as shown in Fig. 5D. Porosity measurements (Fig. 5, E-G) indicate clear differences between patients’ samples at the microscale. Considerable larger pores (red areas in Fig. 5, F and G) are present in patients CF-00017 and CF-00019, compared to the other patients. Moreover, there was an approximately 1.5-fold and 3-fold higher measured porosity in patients CF-00017 and CF-00019 samples compared to CF pooled patient sample and other patients (Fig. 5E), respectively.

**Figure 5.**
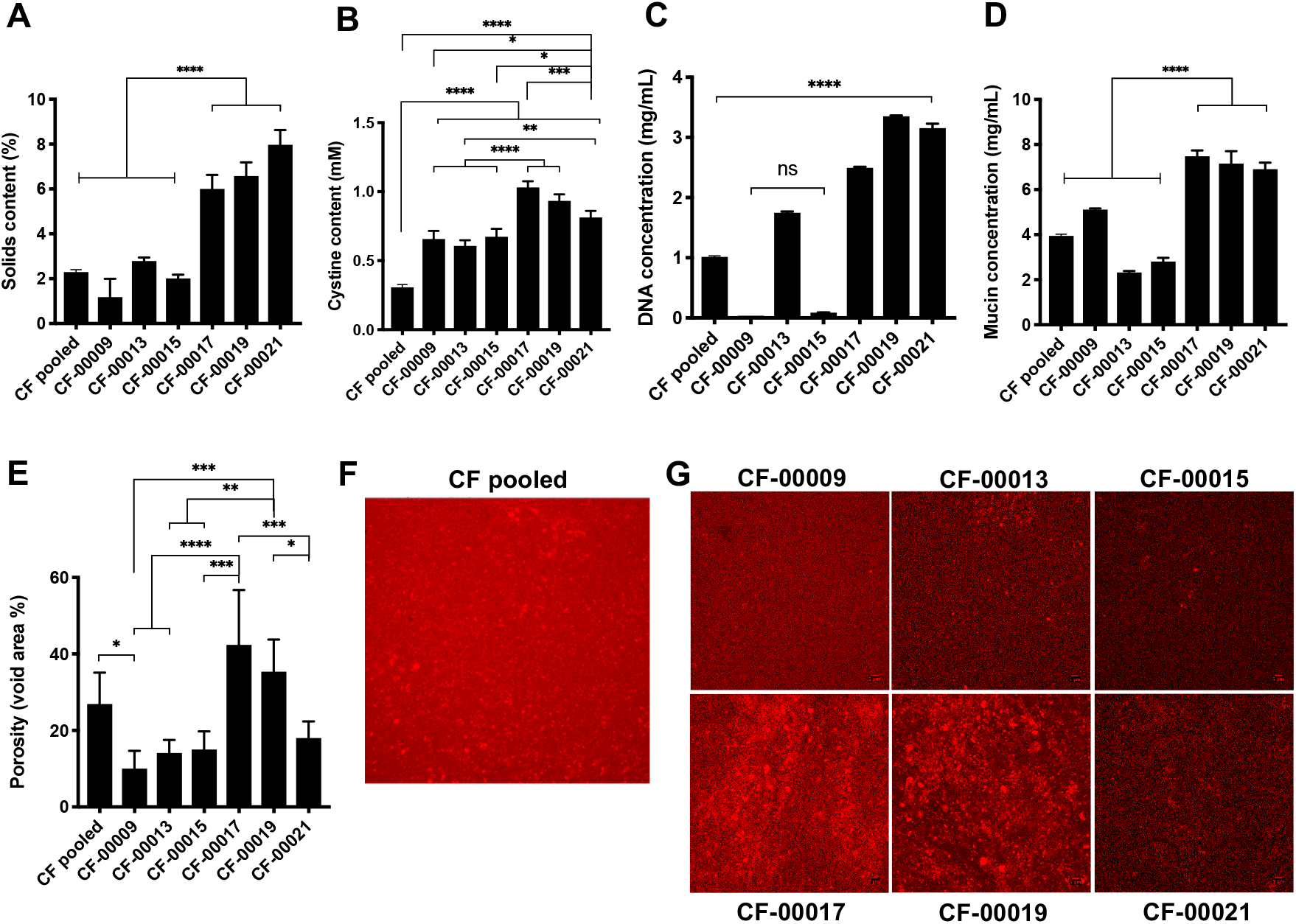
CF sputum samples biochemical composition is significantly different between patients. (A) Solids content, (B) cystine content (i.e. disulfide bonds), (C) DNA concentration, (D) mucin concentration, (E-G) void area (i.e. porosity) measurements. Data represents mean ± SD (n = 3). One-way analysis of variance (ANOVA) with Tukey’s multiple comparisons test.

### Multiple particle tracking in CF sputum

Next, we wanted to measure the diffusivities of our mucus-penetrating peptides out of structural context of phage and compare their transport with poly(ethylene) glycol (PEG) chemistries of similar size scale. To compare our peptides with PEG, here we performed multiple particle tracking experiments with fluorescent carboxyl-modified 100 nm nanoparticles either conjugated with mucus-penetrating peptides or 1 kDa mPEG and measured their transport rates in CF sputum at the microscale. Based on our permeation assays using phage, two patient samples CF-00015 and CF-00021 were chosen for comparison using particle tracking. Fig. 6 AB and Table 2 show the mean squared displacement (MSD) per μm^2^ over time scale τ = 1 s for each patient. As shown in Fig. 6 and Table 2, 100 nm nanoparticles conjugated to PEPTIDE-2 had approximately 1.5-fold better transport rates compared to PEPTIDE-4 functionalized nanoparticles in patient CF-00015. Nanoparticles conjugated with peptides performed up to 1.3-fold better in patient CF-00015 compared with mPEG conjugated nanoparticles, while in patient CF-00021, mPEG exhibited up to 1.5-fold higher MSD per μm^2^ at τ = 1 s. In addition, patient CF-00015 had approximately 50-fold higher MSD values compared to patient CF-00021. This result correlates with phage permeation results, where phage clones permeated approximately 5000-fold higher through patient CF-00015 compared to patient CF-00021 (Fig. 4).

**Figure 6.**
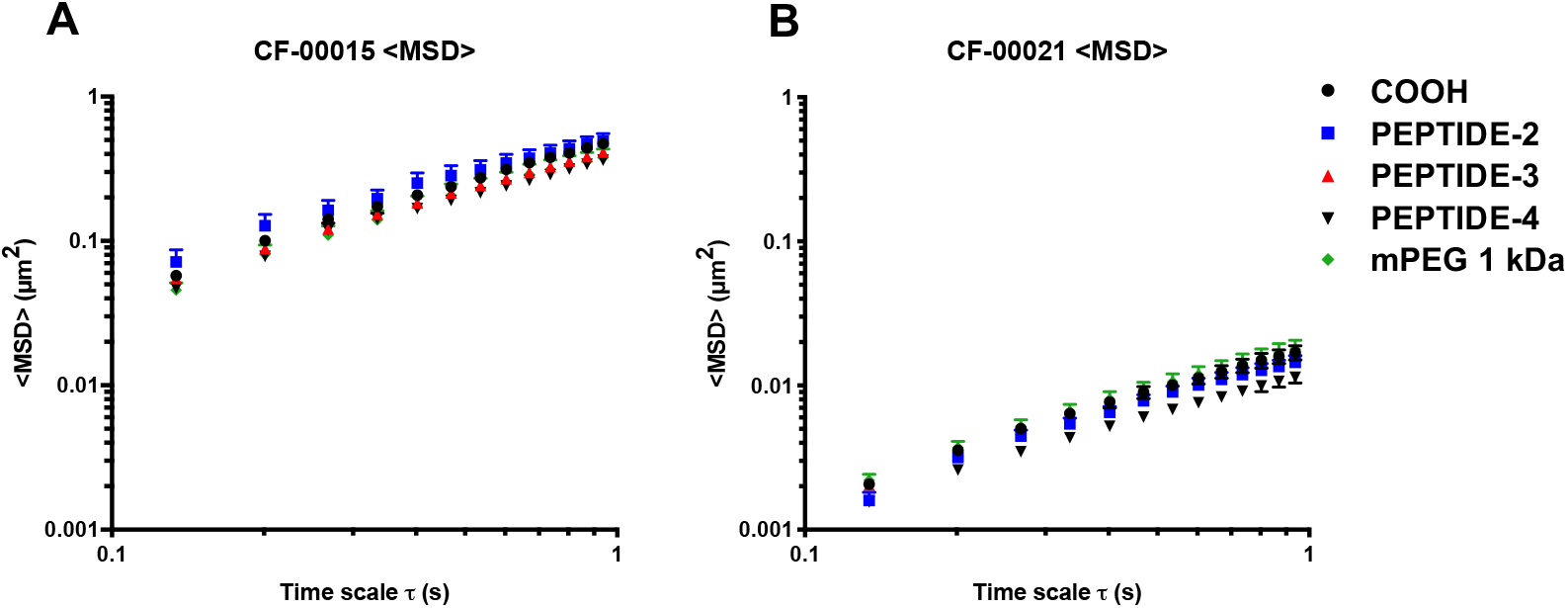
Multiple particle tracking of carboxylated 100 nm fluorescent nanoparticles conjugated to clones 2-4 peptide sequences and mPEG in CF sputum samples from patients (A) CF-00015 (ns, not statistically significant between samples at time scale τ = 1 s) and (B) CF-00021 (p=0.0002 for COOH vs Clone 4, p=0.0015 for Clone 3 vs. Clone 4, and p<0.0001 for Clone 4 vs. mPEG 1K). One-way analysis of variance (ANOVA) with Tukey’s multiple comparisons test. Data represents mean ± SD (n = 3).

**Table 2.** Characterization of fluorescent carboxyl-modified 100 nm nanoparticles conjugated onto their surface via EDC chemistry with clones 2-4 peptides sequences, and mPEG 1KDa measured by DLS at 25 °C. Ensemble mean squared displacement (MSD) per μm^2^ at time scale τ = 1 s for patients CF-00015 and CF-00021. Data represents mean ± SD (n = 3).

| Surface chemistry | Diameter (nm) | Zeta-potential (mV) | PDI | <MSD> ($\mu\text{m}^2$) at time scale $\tau = 1$ s | |
| --- | --- | --- | --- | --- | --- |
|  |  |  |  | CF-00015 | CF-00021 |
| COOH | 110.4 ± 0.6 | -52.3 ± 0.5 | 0.019 | 0.501 ± 0.176 | 0.018 ± 0.002 |
| PEPTIDE-2 | 113.3 ± 0.9 | -49.5 ± 0.3 | 0.014 | 0.517 ± 0.247 | 0.015 ± 0.006 |
| PEPTIDE-3 | 113.8 ± 1.5 | -48.6 ± 0.9 | 0.013 | 0.442 ± 0.104 | 0.017 ± 0.003 |
| PEPTIDE-4 | 113.5 ± 2.8 | -47.2 ± 1.4 | 0.020 | 0.386 ± 0.091 | 0.012 ± 0.004 |
| mPEG 1 kDa | 116.6 ± 2.3 | -45.0 ± 1.1 | 0.015 | 0.399 ± 0.222 | 0.019 ± 0.003 |

### Mucus barrier and cell uptake co-culture assay indicate higher uptake of mucus-penetrating peptide-conjugated nanoparticles

Building on our work to identify and characterize peptides that transport through the CF-like and CF patient sputum, we wanted to confirm their ability as surface chemistries to facilitate intracellular uptake of nanoparticles. Nanoparticles are promising carriers as for transmucosal delivery due to their ability to deliver and protect large amounts of therapeutic cargo while minimizing off-target toxicity. However, these nanoparticles must overcome the transport barriers presented by both the mucus and cell surface. The enhanced transport of nanoparticles across mucus barriers using different surface chemistries, such as hydrophilic, net-neutral polymer coatings, has been extensively demonstrated [7, 17, 20, 23, 57–60]. However, even after penetration through the mucus barrier, nanoparticles must bind and enter cells to deliver their payload. Previous studies have demonstrated that the cellular uptake of nanoparticles depends on their physicochemical properties such as size, shape and surface chemistry [61–64]. While existing surface coatings on nanoparticles improve mucus penetration, there are challenges associated with achieving intracellular uptake in CF-affected cells. Consequently, we confirmed that our mucus penetrating peptides as surface coatings could permeate through CF sputum and achieve cell uptake. Towards this end, we incubated 1 mg/mL of fluorescent nanoparticles functionalized with mucus-penetrating peptides or mPEG in a transwell co-culture with CF sputum on the donor compartment and CuFi-1 bronchial epithelial cells in the receptor compartment for 2 hours at 37°C, and uptake was measured by flow cytometry (Fig. 7A). Prior to incubation in the co-culture setup, the nanoparticles were monodisperse and stable in 1X PBS and in bronchial epithelial growth medium (BEGM) (Supplementary Table 3). From the transwell co-culture assay of nanoparticle uptake, there was approximately 3-fold higher uptake of PEPTIDE-2 conjugated nanoparticles compared to non-modified, carboxylated nanoparticles and mPEG 1KDa conjugated nanoparticles (Fig. 7B). PEPTIDE-4 conjugated nanoparticles exhibited 2.6- and 2.2-fold significantly improved uptake compared to carboxylated nanoparticles and mPEG 1kDa conjugated nanoparticles, respectively (Fig. 7B).

**Figure 7.**
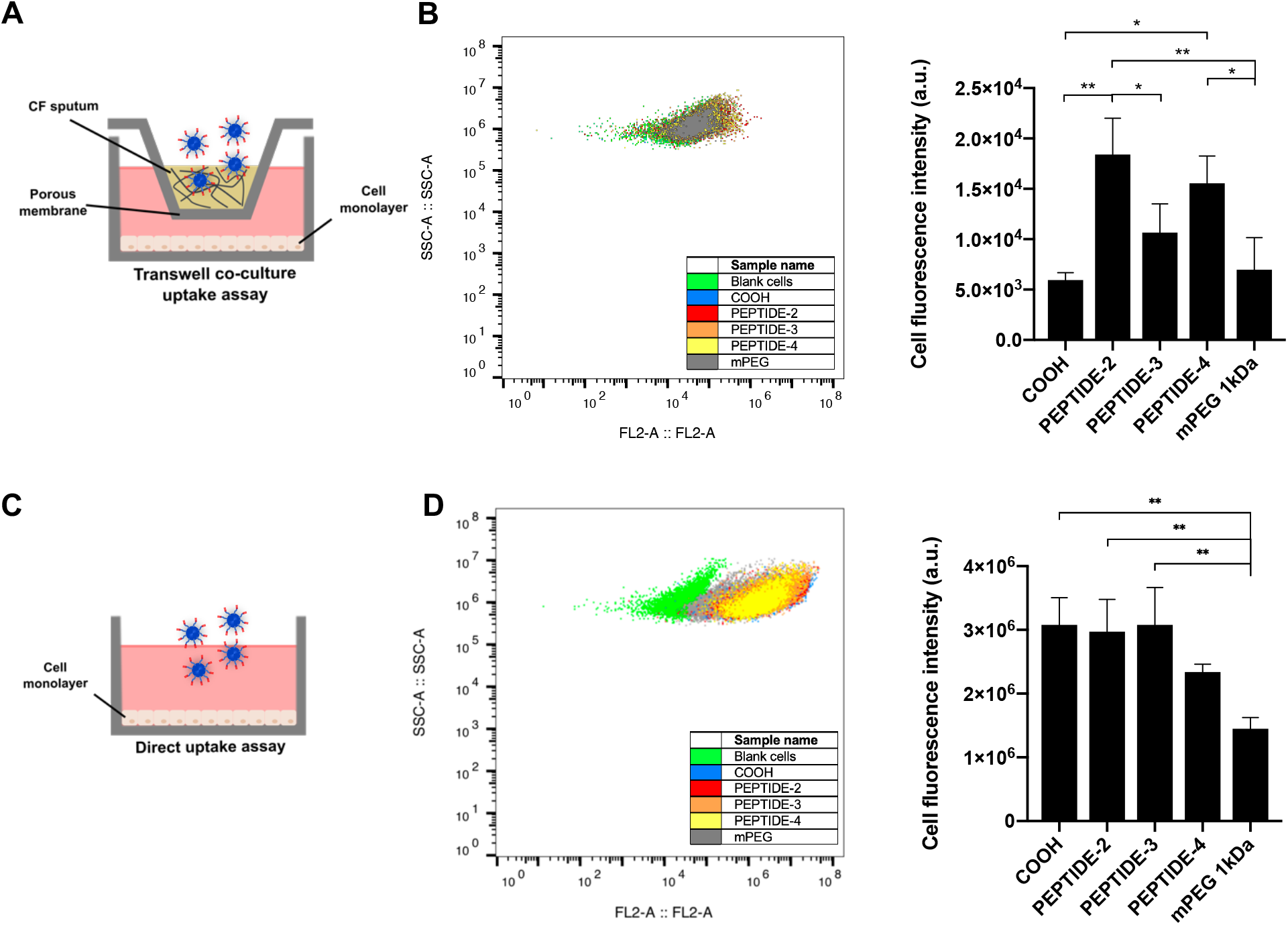
Overview of nanoparticles cell uptake and ranking of the fluorescence intensity of cells. CuFi-1 cells were incubated with 100 nm yellow-green carboxyl-modified nanoparticles conjugated onto their surface via EDC chemistry with peptides sequences 2-4, and mPEG 1kDa for 2 hours at 37°C before flow cytometry measurements. (A-B) transwell co-culture nanoparticle uptake assay with CF sputum and CuFi-1 cells; (C-D) CuFi-1 cells and nanoparticle uptake study. One-way analysis of variance (ANOVA) with Tukey’s multiple comparisons test. Data represents mean ± SD (n = 3).

To confirm that improved uptake was due to cellular uptake and not decreased delivery due to hindered diffusion through CF sputum, we tested the ability of conjugated nanoparticles to enter cells directly without permeation across CF sputum. Here, we incubated the CuFi-1 cells with 25 μg/mL of each fluorescent nanoparticle suspension for 2 hours at 37°C and quantified uptake by flow cytometry (Fig. 7C). As shown in Fig. 7D, CuFi-1 cell uptake of carboxylated, PEPTIDE-2, and PEPTIDE-3 functionalized nanoparticles was approximately 2.1-fold higher compared to 1kDa mPEG conjugated nanoparticles. PEPTIDE-4 conjugated nanoparticles exhibited approximately 1.6-fold higher cell uptake compared to 1 kDa mPEG conjugated nanoparticles.

## Discussion

For successful delivery, drug and gene therapies must overcome poor diffusion through the mucus barrier and after crossing the mucus layer, must also achieve intracellular uptake. To address these challenges, we have developed a phage transport assay to screen T7 phage-displayed peptide libraries in CF mucus to identify mucus-penetrating peptides as surface coatings that facilitate transport through mucus and ultimately, penetrate the affected cells. The T7 phage vector can be genetically engineered to multivalently display 415 copies of a peptide on its gp10 protein coat [65] and has size features (~60 nm diameter) on the scale of conventional synthetic nanoparticles. After repeated screening of T7 phage library (~10^5^ diversity of random peptides) against CF mucus (Fig. 1A), we identified phage able to transport through mucus with more than 200-fold increased output in the third round eluates compared to the first round of screening (Fig. 1B). This increase in output is indicative of enrichment and selection of phage clones with favorable properties [47], in this case for mucus penetration. Previously, using a different M13 phage library against CF mucus-like reconstituted mucin, we identified phage with up to 17.5-fold improved output ratios in mucus in the fourth round of screening, and phage-displayed peptides demonstrated up to 2.6-fold improved transport in mucus compared to wild-type control (i.e. non-recombinant, insertless phage) [66]. However, this low-throughput sample size (20-100 clones) from traditional Sanger sequencing, while providing valuable information, presents a limited repertoire of the complete sequence space (less than 0.01% of available sequences), and as a result, it is likely that better performing peptides are not identified [53]. Second, the low copy display of peptides on the p3 coat of M13 (i.e. 5 copies versus 415 for T7) has a small surface area and does not allow for multiple interactions with the mucus substrate, which is needed to identify peptides that have desired weak but reversible interactions with mucus needed for penetrating transport [67]. Finally, while previous work focused on phage transport through reconstituted mucin, it is necessary to validate transport in sputum samples of CF patients; building insights from patient samples advances translational aspects in therapeutic applications of the mucus- and cell-penetrating peptide carriers.

We utilized a bioinformatics workflow to count and sort the frequencies of individual phage-displayed peptides in each eluate from the rounds of screening against CF mucus. A large diversity of peptides was observed in the output eluates in the first-round replicates (Supplementary Table 2), possibly due to the complexity of the sample. However, there were fewer unique peptides after each round, with a ~12- and 15-fold decrease after the second and third rounds compared to the first round, respectively (Supplementary Table 2). This observation is in agreement with the expected process of selection, where there is an increase in specificity of phage-displayed peptides over additional rounds of selection and a decrease in their sequence diversity [47, 54]. These findings suggest that mucus acts as a selection pressure for unique mucus-penetrating peptides after each round of screening. From selection, a small set of mostly hydrophilic and slightly negative or close to neutral net-charge peptide sequences were selected for subsequent transport studies in patient samples (Table 1). Of interest, three of the four discovered peptides presented homology to human mucin proteins in a basic local alignment search tool (BLASTp) database search. Sequence homology of peptide sequences to their targets have been previously reported in phage display biopanning experiments in the human vasculature, collagen, and peanut allergens, among others [68–73]. It is feasible that mucus biomimetic peptides might shield or have less hindered intermolecular interactions with mucins [74], enabling their diffusion across mucus, although the selected peptide sequences are small in size and may not represent a statistically significant alignment with proteins from BLASTp databases.

The combination of phage display libraries with high-throughput sequencing provided an in-depth study of the phage selection process and also the identification of the most abundant peptides sequences in each round of screenings. We determined the overall amino acid composition of the peptides from each round of selection (Fig. 3). There was an increase in glycine, serine, and acidic residues, such as glutamic acid and aspartic acid, and a decrease in hydrophobic and basic residues, such as histidine, lysine, and arginine. The changes in amino acid composition during selection can be attributed due to intermolecular interactions (or lack thereof) between peptide substrates with mucins present in mucus. Mucins, the main non-aqueous component of mucus, are heavily glycosylated proteins with an overall net-negative charge due to high sialic acid and sulfate content [75]. Subsequently, it has been demonstrated that positively-charged molecules typically exhibit low permeability across the mucus barrier due to interactions with the glycoproteins and lipids in the mucus [9, 76]. Furthermore, the diffusion of hydrophobic molecules in mucus is hindered compared to hydrophilic molecules due to adhesive interactions with hydrophobic domains stabilized by disulfide bonds present in mucins [77]. These findings suggest that hydrophobicity and charge can influence diffusion in mucus due to intermolecular interactions with mucins. Moreover, we calculated the physicochemical properties (i.e. charge and hydrophobicity) of the most twenty abundant sequences in each round of screening, and the *in silico* analysis indicated that the peptide pool presented mostly negative net-charge and hydrophilic sequences over rounds of selection. The prevalence of hydrophilic residues is consistent with prior work where hydrophilic polymers provided an inert surface for mucus, or “muco-inert”, which minimizes mucin interactions for improved particle transport [17].

We confirmed that identified phage-displayed peptides improved diffusion of phage in reconstituted CF mucus and sputum from CF patients (Fig. 5). We pooled samples from patients to minimize patient-to-patient variability and were able to demonstrate up to ~600-fold greater permeation of the phage-displayed peptides in CF sputum, compared to the positively charged control phage clone. Since mucus has an overall negative net-charge due to the presence of carboxyl or sulfate groups on the mucin proteoglycans, it is expected that neutral to negatively charged molecules diffuse faster compared to positively charged molecules due to lack of strong electrostatic-driven binding to mucin fibers [57]. Indeed, previous studies corroborated that neutral and negatively charged nanoparticles have demonstrated higher diffusivities in mucus compared to positively charged particles [23, 78].

We also studied transport of the identified phage-displayed peptides in sputum samples from individual CF patients. Transport through samples was dependent of the heterogeneity of mucus. It has been reported that increased solids, mucin, DNA, and disulfide bonds content reduced pore size in the sputum mesh [52, 60, 79]. Surprisingly, patients 17 and 19 showed higher amounts of solids, disulfide bonds, DNA, and mucin content, yet presented larger porosities. Those patients presented highest permeabilities of the tested phage clones. Patient 15, in contrast, presented lower solids, disulfide bonds, DNA, and mucin content, while presenting lower porosity measurements but still higher phage permeation. Patient 21 presented a greater barrier for phage permeation, which can be explained by the combination between higher solids, disulfide bonds, DNA, and mucin content with low porosities, imposing a challenging barrier for particles and molecules to diffuse through. These findings suggest that the properties governing diffusion of particles in sputum at the microscale are patient-dependent and not controlled only by the mucus composition alone. The permeation of molecules and particles in sputum are most likely governed by a combination of factors, including mesh size, mucin concentration, and physical entanglements between mucin fibers, DNA, and disulfide cross-linking [79]. Patient heterogeneity for CF samples has been previously reported in the literature, with sample variations even between the same patient intra-day, or within the same sample [80], which suggests that CF sputum properties at the microscale are patient-specific. Our findings motivate future studies to screen phage-displayed peptide libraries against individual CF patients for personalized medicine strategies.

To query the transport of the identified peptides in the CF sputum microenvironment, we conjugated synthetic peptides to carboxyl-modified polystyrene nanoparticles of 100 nm diameter and compared their mean squared displacement (MSD) in patient sputum with nanoparticles conjugated to mPEG 1 kDa. Recent studies have shown hydrophilic, net-neutral charge PEG polymer and polymer conjugates can improve transport of particles in CF sputum up to 90-fold compared to uncoated particles and improve therapeutic activity of antibiotics such as tobramycin against bacterial biofilms commonly associated with CF infections [17, 81–83]. Here, we used PEG 1 kDa as it is expected that similar molecular weights among peptides and PEG conjugates would experience similar steric hindrance within the mucin fiber mesh network, and potential differences in diffusivities would be mostly attributed to electrostatic and hydrophobic interactions with mucus components. Interestingly, we found that the diffusion of carboxylated uncoated nanoparticles was close or sometimes better than the conjugated chemistries in both patients. This could be partially attributed to the reduced size compared to the conjugated nanoparticles, since those were uncoated particles. While the addition of peptides and mPEG to carboxylated nanoparticles slightly increased the zeta-potential, the overall net-charge of all the particles remained negatively charged, which could affect diffusion of particles in CF sputum, as previously described. Nevertheless, our findings are in agreement with previous reports in the literature where 40% of uncoated carboxyl-modified polystyrene nanoparticles were able to diffuse in respiratory mucus; those results suggest that particles might be less adhesive to respiratory mucus than they are to cervical mucus, possibly due to interactions with endogenous surfactants present in the airway mucus [20].

To measure transport of our peptide conjugated nanoparticles in individual representative patients, we selected two patients that presented similar porosities (Fig. 7E). However, patient 21 showed much higher solids, DNA, disulfide bonds, and mucin content compared to patient 15, which might partially explain such differences in the MSD results. The probability of potential adhesive interactions between the tested 100 nm nanoparticles with components in sputum via electrostatic and hydrophobic interactions might be augmented in patient 21 due to higher concentrations of its components. Moreover, recent studies have found that decreased diffusivity of 100 nm PEGylated nanoparticles in CF sputum correlated with decreased measured forced expiratory volume after 1 second (FEV_1_), the primary measurement for pulmonary function and lung disease severity in CF [79]. These data suggest that sputum may function as a greater barrier to drug delivery systems in some patients compared to others. Nevertheless, we conclude that the results obtained with multiple particle tracking correlate well with the results from bulk diffusion in those individual patients tested. With patient 15, clone 2 peptide showed best transport compared to carboxylated, PEGylated, and other peptides conjugated to nanoparticles, whereas in patient 21, all tested nanoparticles exhibited transport hindered by the CF sputum.

After mucus penetration, it is highly desirable that transmucosal drug delivery systems can achieve cell penetration in order to deliver their therapeutic payload intracellularly. Cell-penetrating peptides have recently demonstrated the potential to enhance the mucosal delivery of drugs and particles [84, 85]; however these positively charged peptides form strong interactions with the negatively charged mucin glycoproteins, thereby hindering transport and actually, preventing their ability to achieve cell-penetration [86]. Furthermore, while promising, net-neutral charge polymers and polymer conjugates demonstrate improved mucus transport, they may face hindered cellular internalization and escape [16, 33–37, 87–89], thereby dampening the enthusiasm for their therapeutic use. Hence, we confirmed the ability of our peptides to achieve both mucus penetration and cellular internalization by measuring transport of the identified mucus-penetrating peptides conjugated to 100 nm nanoparticles in a co-culture assay of CF sputum and lung epithelial cells possessing the prevalent cystic fibrosis homozygous delta F508 mutation. In CF, the three basepair deletion of codon 508 (known as F508del) results in a loss of a phenylalanine residues on chromosome 7 and is the most common mutation amongst several identified CFTR mutations [90], with approximately 70% prevalence worldwide [91]. From our transport studies, we found that select mucus-penetrating peptides not only improved the bulk diffusion of phage in CF sputum from patients ~600-fold more than a positively charged control phage clone but also achieved up to 3-fold significantly higher cell uptake of conjugated nanoparticles compared to non-modified carboxylated- and mPEG-conjugated nanoparticles. We also found that the PEG coating significantly decreased cell uptake of nanoparticles by up to 2-fold compared to mucus-penetrating peptides- and noncoated carboxylated nanoparticles. Collectively, these results indicate that the identified hydrophilic, net-neutral charge peptides that have sequence similarity to mucins successfully traverse the complex CF sputum barrier and improve intracellular uptake of conjugated nanoparticles to lung epithelial cells.

## Conclusions

Our findings demonstrate that using phage display libraries with high throughput sequencing may be an effective strategy to identify mucus-penetrating peptides with improved transport across CF sputum from patients and promote cell internalization of drug delivery systems. These peptides may effectively serve as conjugates or surface modifications to potentially increase the amount of drug and drug carriers delivered to the CFTR-mutated epithelia to improve clinical outcomes. Moreover, the combination regimen using different peptide-conjugated drug carriers could be used as an alternative to reduce potential immunogenic responses over long-term, repeated administration of therapeutics [28, 92]. Screening of peptide-presenting phage libraries against samples from individual patients could be used to identify patient-specific mucus- and cell-penetrating peptides to explore personalized medicine applications. While this work focuses on mucus environment in CF, the use of mucus-penetrating peptides can be extended to develop the next generation of therapeutics to advance gene and drug delivery through various mucosal barriers, including the lung airways, eyes, gastrointestinal tract, and cervicovaginal tract.

## Materials and Methods

### Library construction

A combinatorial library of heptapeptides flanked by a pair of cysteine residues displayed on T7Select415-1 phage (Novagen, WI) was constructed according to the manufacturer’s protocol. Briefly, degenerate NNK-oligonucleotides (Supplementary Table 1) encoding a library of heptapeptides with the general structure CX7C, were synthesized and obtained from Integrated DNA Technologies (IDT, IL). To create the library insert, oligonucleotides (LIBF and LIBR, Supplementary Table 1) were annealed, extended with DNA polymerase I Klenow fragment (NEB, MA), double digested with HindIII-HF/EcoRI-HF (NEB, MA), and cloned into the T7Select 415-1 HindIII/EcoRI vector arms to enable peptide display on 415 copies of the T7 gp10A phage capsid protein as a C-terminal fusion.

### Phage selection in CF mucus

The generated T7 phage CX7C library was iteratively screened against a cystic fibrosis mucus model (CF mucus model) in a 3.0 μm polyester membrane 24-well transwell system (Corning, MA) to identify mucus-penetrating peptides. The CF mucus model was prepared according to literature reports of composition to model sputum of CF patients. The model is an aqueous mixture of 40 mg/mL mucin from porcine stomach type III (Sigma Aldrich, MO) [93], 1.4 mg/mL herring sperm DNA (Promega, WI) [94], 5 mg/mL egg yolk from chicken (Sigma Aldrich, MO) [95], 0.9 mg/mL lactoferrin human [95], 5 mg/mL sodium chloride, and 2.2 mg/mL potassium chloride to give the concentration of ions found in CF sputum [95]. The solution was adjusted with 0.5 M hydrogen chloride to pH 6.5, which is the estimated pH of CF airway mucus [96]. The reagents were transferred to a sterile tube, mixed for 2 h in a tube rotator at 5 rpm (Thermo Fisher Scientific, MA), and used within 4 h.

To set up the transwell system, the basolateral compartment (receiver) in the transwell was filled with 600 μL of phosphate buffered saline (PBS), and the apical compartment (donor) was filled with 100 μL of CF mucus. For the first round of selection, an initial phage amount of 4.2 x 10^9^ phage plaque forming units (pfu) was added on top of the mucus layer in the donor compartment and incubated for 1 h at room temperature (25°C). At 5, 10, 15, 30, and 45 minutes, 10 μL aliquots of the eluates were taken for further quantification. After 1 h, the entire eluate was collected from the basolateral side and titered using standard double-layer plaque assay to quantify phage concentration. The eluted phage library was amplified in BL21 E. coli (Novagen, WI) to make more copies, which was quantified by plaque assay prior to the next round of selection. Two additional rounds of selection were performed with initial amounts of 5.2 x 10^6^ pfu and 4.2 x 10^9^ pfu, respectively.

### Next-generation sequencing of phage libraries selected against CF mucus

#### Library preparation for next-generation sequencing

After the third round of selection against the CF mucus model, the eluates from 15, 30 and 60 minutes from the three rounds, and two replicates each were amplified in BL21 E. coli (Novagen, WI). In addition, the naïve library (i.e. starting library) was amplified over three rounds without selection for further use as controls to account for bias due to amplification growth. Next, the amplified phage eluates were prepared for next-generation sequencing following a library preparation workflow to the manufacturer’s protocol (Illumina, CA); the naïve library was also similarly prepared in order to determine the diversity of the CX7C library (i.e. how many unique clones). Samples were heat denatured at 100°C for 15 minutes in a heated dry bath (Thermo Fisher Scientific, MA), and their DNA was amplified by PCR using 2x KAPA HiFi HotStart ReadyMix (KAPA Biosystems, MA) with 2.5 μL of phage DNA template and 10 μL of 1μM primers (NGS1F and NGS1R, see Supplementary Table 1). The PCR conditions were carried out as recommended by the manufacturer. The obtained PCR amplicons were purified with AMPure XP beads (Beckman Coulter, IN), and size was confirmed by gel electrophoresis in a 2% agarose gel. Next, sequencing adaptors and unique dual barcodes combinations were attached to each library PCR amplicons using the Nextera XT Index Kit primers (index 1 N7XX and index 2 S5XX, Illumina, CA), and the amplicon size and concentration were confirmed using a Bioanalyzer (Agilent) and Nanodrop (Thermo Fisher Scientific, MA), respectively. Samples were diluted to the final concentration of 40 nM in 10 mM Tris buffer, pH 8.5. One microliter aliquots of each sample diluted DNA with unique indices (i.e. barcodes) were pooled together and submitted for sequencing to The University of Texas at Austin GSAF sequencing facility on a MiSeq system to run single reads of 300 nucleotides.

#### Translating sequencing reads to peptides and frequency counts

Sequences were automatically demultiplexed by MiSeq Reporter software and the obtained FASTQ files were analyzed by FastQC version 0.11.5 (http://www.bioinformatics.babraham.ac.uk/projects/fastqc/) for quality control checks on raw sequence data. Next, sequences were converted from fastq to fasta format using FASTX-Toolkit fastq_to_fasta tool version 0.0.13.2 (http://hannonlab.cshl.edu/fastx_toolkit/index.html), and the resulting fasta files were processed using a custom Bash script to find the variable library region by aligning to sequences flanking both sides of the displayed CX7C peptide (available under request). For each sample, subsequences corresponding to the phage display variable region (including the C-C constrained library insert) were transcribed and translated into amino acids using a custom Python script, version 2.7.12 (available under request). Then, sequence reads encoding unique peptide sequences had their frequency count calculated and sorted according to their abundance in each sample. An asterisk (*) indicates the presence of a stop codon. Sequences were disregarded in subsequent analysis if they contained a stop codon in the first three amino acid positions in the variable region. An overview of the number of reads that passed each processing step can be found in Supplementary Table 2. For each sample, the top 20 most abundant peptide sequences were compiled, and their physicochemical properties were determined *in silico* using Protein Calculator v3.4 (The Scripps Research Institute, La Jolla, CA), and the grand average of hydropathy (GRAVY), a score indicative of the hydrophobicity of a sequence, was calculated according to Kyte-Doolittle [55]. Sequence database analysis was performed using Basic Local Alignment Search Tool (BLASTp, U.S. National Library of Medicine, Bethesda, MD). In addition, the selected peptide sequences were compared to databases encompassing previously isolated peptides known as “target-unrelated peptides” (TUP) motifs [97, 98]; peptide sequences that appeared in the search were excluded from subsequent analysis. These databases have identified a large set of sequences with undesired proliferation advantages and false positive target-unrelated peptides.

#### Cloning of selected sequences into T7 phage vector

Oligonucleotides encoding the peptides of interest were designed and obtained from IDT (Supplementary Table 1). Primers (LIBF and respective CLNX, Supplementary Table 1) were annealed, extended with DNA polymerase I Klenow fragment (NEB, MA), double digested with HindIII-HF/EcoRI-HF (NEB, MA), and cloned into the T7Select 415-1 HindIII/EcoRI vector arms according to the manufacturer’s protocol (Novagen, WI). Individual phage plaques (i.e. areas of cell lysis on overlaid agar plates indicative of phage with individual cloned sequences) were selected from titer plates and grown in liquid culture of BL21 E. coli (Novagen, WI) to amplify the amount of phage. To confirm cloning, DNA of the individual clones were PCR amplified for 20 cycles following the manufacturer’s temperature settings with T7 primers (T7F and T7R, Supplementary Table 1), and sequences encoding for selected peptides were confirmed by Sanger DNA sequencing.

### CF sputum sample collection

Spontaneously expectorated sputum samples were collected from patients at the CF clinic at Dell Children’s Medical Center of Central Texas Cystic Fibrosis Center, Dell Children’s Pulmonology Clinic, Seton Healthcare Family (n = 21) following protocols approved by the Institutional Review Board of the University of Texas at Austin under study number 2016-03-0104. Informed consent was obtained from all study participants. Samples were stored at −80°C immediately after collection and thawed on ice for subsequent experiments. Age, gender, and any inhaled medications taken by the patients were recorded. Sputum samples with visible amounts of saliva were excluded from experiments.

### Biochemical characterization of CF sputum

The sputum of CF patients was characterized for solids content, mucin concentration, DNA concentration, and cystine content. Aliquots of individual sputum samples were pretreated with 20 U/mL of DNase I (NEB, MA) for subsequent fluorometric assays of mucin and DNA concentrations. Mucin concentration was determined by the reaction of 2-cyanoacetamide (Sigma Aldrich, MO) with O-linked glycoproteins of the mucin, as previously described [20, 79]. Briefly, sputum aliquots were diluted 20-fold and homogenized by vortexing for at least 30 minutes. Next, 50 μL of diluted sputum were mixed with 60 μL of an alkaline solution of 2-cyanoacetamide (previously prepared with 200 μL of 0.6 M 2-cyanoacetamide and 1 mL of 0.15 M NaOH) and incubated at 100°C for 30 minutes. To stop the reaction, 0.5 mL of 0.6 M borate buffer, pH 8.0, was added to the mixture. A standard curve was generated using known concentrations of mucin from bovine submaxillary gland (Sigma Aldrich, MO). Fluorescence intensity was measured at excitation and emission wavelengths of 340 and 420 nm (Infinite M200, Tecan, Switzerland), respectively.

DNA concentration of patient sputum was determined by the reaction of 3,5-diaminobenzoic acid dihydrochloride (DABA; Sigma Aldrich, MO) with DNA aldehydes, as previously described [20, 79]. Briefly, sputum aliquots were diluted 5-fold and homogenized by vortexing for at least 30 minutes. Next, 30 μL of diluted sputum were mixed with 30 μL of a 20% (w/v) DABA solution and incubated at 60°C for 1 hour. To stop the reaction, 1 mL of 1.75 M HCl was added to the mixture. A standard curve was generated using known concentrations of DNA from herring sperm (Promega, WI). Fluorescence intensity was measured at excitation and emission wavelengths of 400 and 520 nm (Infinite M200, Tecan, Switzerland), respectively.

The concentration of mucin disulfide bonds (i.e., cystine) was determined as previously described [79, 99]. Briefly, 50 μL sputum aliquots were centrifuged at 21,000 x g for 1.5 hours, and the supernatant was removed. Next, sputum aliquots were diluted 10-fold the original volume in 8 M guanidine-HCl (Sigma Aldrich, MO). Samples were pretreated with 10% (v/v) 500 mM iodoacetamide (Sigma Aldrich, MO) solution and incubated at room temperature for 1 hour. Then, samples were incubated with 10% (v/v) of 1 M DTT (Sigma Aldrich, MO) at 37°C for 2 hours to quench the excessive iodoacetamide and reduce all the preexisting disulfide bonds. Samples were buffer-exchanged with 50 mM Tris-HCl, pH 8.0 (Thermo Fisher Scientific, MA) in 7K MWCO Zeba desalting columns (Thermo Fisher Scientific, MA) to remove excess reagents. A standard curve was generated using known concentrations of L-cysteine (Sigma Aldrich, MO) in 50 mM Tris-HCl, pH 8.0. One hundred microliters of samples were added to a black 96-well plate with equal volume of 2 mM monobromobimane (Sigma Aldrich, MO) in 50 mM Tris-HCl. After 15 min incubation at room temperature, fluorescence intensity was measured at excitation and emission wavelengths of 395 and 490 nm (Infinite M200, Tecan, Switzerland), respectively.

The solids content of sputum was determined by freeze-drying. Briefly, sputum samples were frozen in liquid nitrogen and placed in a lyophilizer (Labconco, MO) for 24 hours. Samples were weighed before and after lyophilization, and their solids mass ratio was determined as the percent solids content. The data presented is the average of three independent experiments.

### Fluorescent dextran staining of CF sputum and porosity measurements

The void area in CF sputum (i.e., porosity) was determined as previously described, with slight modifications [79]. Briefly, 20 μL sputum aliquots were added to custom microscopy chambers prepared using 13-mm diameter adhesive seal spacers (Grace Bio-Labs, OR). Next, 0.3 μl of a 10 mg/mL stock solution of 3kDa Texas red-labeled dextran (Invitrogen, CA) were added to the chamber and mechanically mixed with a pipette tip to distribute it throughout the sample. The chamber was sealed with a coverslip and equilibrated at room temperature for 1 h before imaging. CF sputum samples were imaged using a Zeiss LSM 710 confocal microscope at 63x magnification with image resolution of 105 nm/pixel. The area of the dextran-filled void space (VD) was measured using ImageJ and used to determine the porosity, which was calculated as ϕ = VD/VT, where VT is total area of the image (140 μm × 140 μm).

### Bulk diffusion of T7 phage clones

Diffusion studies of phage-presenting peptide clones were performed in a 3.0 μm polyester membrane 24-well transwell across CF mucus or CF sputum from patients at room temperature (25°C). Briefly, 100 μL of CF mucus or CF sputum was transferred to the donor compartment in the transwell system containing 0.6 mL PBS in the receiver compartment. An initial phage concentration of 2*10^9^ genomic copies (gc) was added to the donor compartment on top of the CF mucus or CF sputum layer. At 15, 30, 45, 60, and 120 minutes, the entire eluate was collected from the receiver compartment and replenished with fresh PBS. Collected eluates were quantified by quantitative PCR (qPCR) as previously described [100] to determine phage concentration (in genomic copies (gc)) using the forward and reverse qPCR primers (Supplementary Table 1). Standard curves were generated with serial dilutions of T7Select 415-1b packaging control DNA for each run. The data presented is the average of three independent experiments.

### Functionalization of nanoparticles with mucus-penetrating peptides

Red or yellow-green fluorescent 100-nm diameter carboxylate-modified polystyrene (PS) nanoparticles (Thermo Fisher Scientific, MA) were conjugated to methoxy-PEG-amine 1KDa (Alfa Aesar, MA) or custom synthesized peptides (LifeTein, NJ) via covalent attachment to COOH groups on the particles via standard EDC chemistry using 1-ethyl-3-(3-dimethylaminopropyl) carbodiimide hydrochloride (Thermo Fisher Scientific, MA) and N-hydroxysulfosuccinimide sodium salt (Thermo Fisher Scientific, MA) in MES buffer (pH 6.0), according to the manufacturer protocol. The molar ratio for conjugation between COOH groups and mPEG-amine 1KDa or peptides was 1:5. After conjugation, nanoparticles were dialyzed for 24 hours against 1X PBS using 100 kDa MWCO Float-A-Lyzer G2 dialysis membranes (Spectrum Labs, MA). Freshly prepared nanoparticle dispersions were characterized by dynamic light scattering (DLS) (Zetasizer Nano, Malvern Instruments, MA). Size and zeta potential measurements were performed in water or PBS at 25°C and BEGM cell culture media at 37°C, at a scattering angle of 173°.

### Multiple particle tracking (MPT) in CF sputum from patients

Fluorescent video-rate microscopy experiments were performed to track the motion of nanoparticles in CF sputum. Thirty-microliter sputum aliquots were withdrawn using a Wiretrol (Drummond Scientific Company) and added to custom microscopy chambers. Next, 1.0 μl of nanoparticles were added to the slide chamber and were mechanically mixed with a pipette tip to distribute the particles within the sample. The chamber was sealed with a coverslip and equilibrated at room temperature for 1 h before imaging. Movies were recorded at room temperature at a temporal resolution of 67 ms for 20 s using an Olympus IX-83 inverted fluorescence microscope with a 100x/1.3 NA oil-immersion objective having an image resolution of 64 nm/pixel and a Hamamatsu Orca Flash 4.0 Plus SCMOS digital camera. For each sputum sample, five videos were collected. Data presented are the average of three independent experiments.

To quantify displacement of particles from real-time imaging, MPT analysis was performed using a custom-written script in MATLAB (Mathworks) based on a previously developed algorithm and detailed software package [101, 102]. Briefly, fluorescent nanoparticles had their centroids and x and y positions determined in sequential images with an intensity threshold. Background noise particles above this threshold were discarded based on particles eccentricity. The time-averaged mean squared displacement <MSD(τ)> was calculated as previously described [23, 79, 103, 104], based on particle displacement as <Δr^2^(τ)> = <[x(t + τ) - x(t)]^2^ + [y(t + τ) - y(t)]^2^>, where τ is time scale between frames.

### Cell culture

CuFi-1 cells (ATCC CRL-4013) were maintained as monolayer cultures in flasks pre-coated with 60 μg/mL solution of human placental collagen type IV (Sigma Aldrich, MO) and grown in bronchial epithelial growth medium (BEGM) supplemented with SingleQuot additives from Lonza (BEGM Bullet Kit, reference CC-3170) and 50 μg/mL G-418 at 37°C and 5% CO_2_.

### Transwell co-culture cell uptake assay

A co-culture diffusion experiment with CF sputum from patients and CuFi-1 cells was performed in 3.0 μm polyester membrane 24-well transwells at 37°C and 5% CO_2_. Prior to co-culture, cells were seeded in the receiver compartment of a transwell at a cell density of 15,000 cells/cm^2^ and grown for 24 hours. Next, 100 μL of CF sputum was transferred to the donor compartment in the transwell system, with 0.6 mL BEGM cell culture media and the CuFI-1 cell monolayer in the receiver compartment. Nanoparticle suspensions were added to the donor compartment on top of the CF sputum layer at an initial concentration of 1 mg/mL. After 2 hours, the transwells and cell culture media were removed, and cells were washed 3 times with ice-cold 1X PBS. To detach the cells, 150 μL of 0.25% trypsin-EDTA was added to each well and incubated at 37°C for 10 minutes. Next, 150 μL of ice-cold 1% FBS in Dulbecco’s phosphate buffered saline was added and cells were transferred to centrifuge tubes. Cells were spun at 125 x g for 5 to 10 minutes and the supernatant was discarded. Cells were resuspended in 250 μL 1X PBS with 1.25 μL 1 mg/mL propidium iodide (PI). Cell uptake of nanoparticles was evaluated by flow cytometry (Accuri, Becton-Dickinson, CA). Data presented are the average of three independent experiments.

### Cell uptake of nanoparticles

Prior to cell uptake experiments, 24-well plates were seeded with CuFi-1 cells at a cell density of 15,000 cells/cm^2^ and grown for 24 hours at 37°C and 5% CO_2_. On the day of experiment, 25 μg/mL suspension of fluorescent nanoparticles was added to cells in 0.5 mL BEGM cell culture media. After 2 hours, the cell culture media was removed, and cells were washed 3 times with ice-cold 1X PBS. To detach the cells, 150 μL of 0.25% trypsin-EDTA solution was added to each well and incubated at 37°C for 10 minutes. Next, 150 μL of ice-cold 1% FBS in Dulbecco’s phosphate buffered saline was added, and cells were transferred to centrifuge tubes. Cells were spun at 125 x g for 5 to 10 minutes and the supernatant was discarded. Cells were resuspended in 250 μL 1X PBS with 1.25 μL PI. Cell uptake of nanoparticles was analyzed by flow cytometry.

### Statistical analysis

Unless otherwise indicated, data are presented as mean ± standard deviation. All statistical analyses were performed using GraphPad Prism 8 (GraphPad Software, La Jolla, CA), at a significance level of p ≤ 0.05, and are indicated in each figure legend.

## Author contributions

JL, HDCS, and DG designed experiments. JL performed experiments. XL, RPM, and XP assisted in performing experiments. DA and DW collaborated with the development of bioinformatics pipelines and assisted with analysis of the deep sequencing data. SHS, JJF and BCM collected patient samples and managed clinical data. JL, HDCS, and DG analyzed the experiments. JL, HDCS, and DG wrote the manuscript.

## Acknowledgements

Research reported in this publication was supported by NIH National Heart, Lung and Blood Institute (NHLBI) of the National Institutes of Health under award number R01HL138251, PhRMA Foundation Research Starter Grant, startup funds from the College of Pharmacy, University of Texas at Austin, the Williams and McGinity Graduate Fellowship from the College of Pharmacy, University of Texas at Austin, and the University Graduate Continuing Fellowship, The Graduate School, University of Texas at Austin. The content is solely the responsibility of the authors and does not necessarily represent the official views of the National Institutes of Health.

## Data availability

Next-generation sequencing data will be made available via the Sequence Read Archive (SRA) under Bioproject number PRJNA527341, submission ID SUB5326131.

## Supplementary Information

### 1. SI Figures

**Figure S1.**
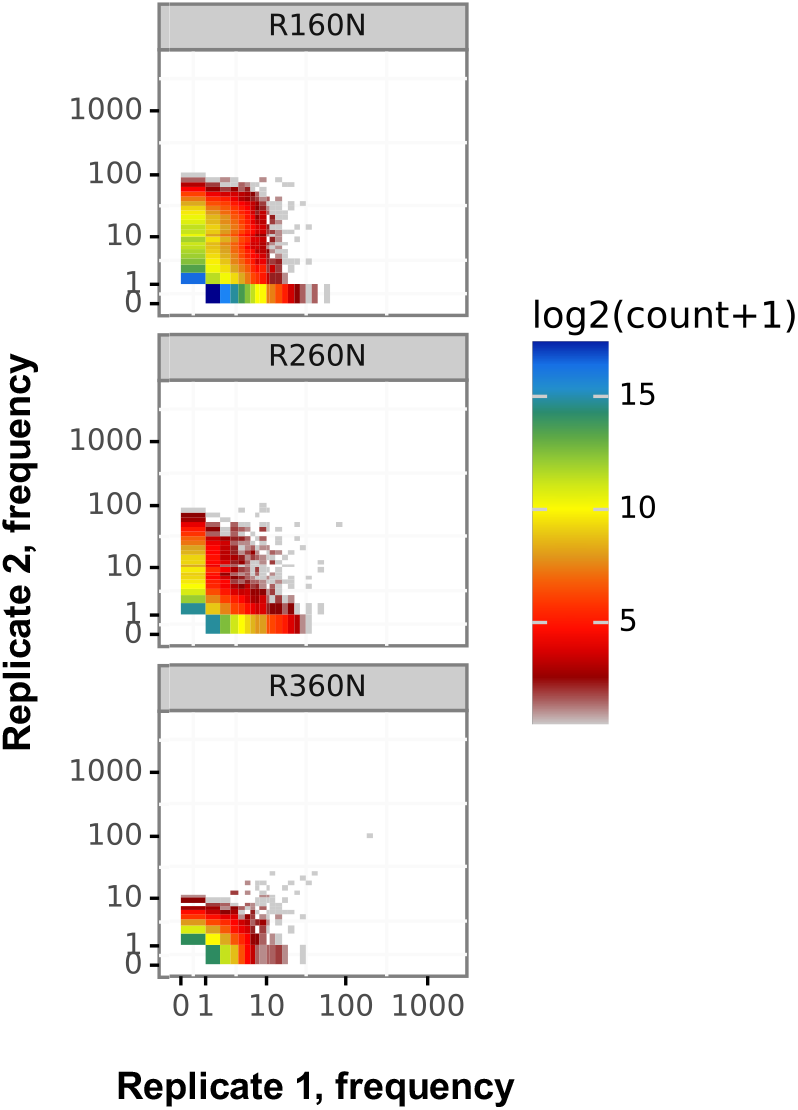
Peptide frequencies over rounds of selection in 60 minutes eluates between two replicates; each point represents a peptide sequence. Heat map represents peptide sequences diversity (i.e. number of unique sequences). The frequencies of specific peptide sequences are reproducible between replicates over three rounds of selection and indicate a convergence of most frequent peptides over rounds, suggesting an enrichment for unique peptide sequences with desired properties (i.e. mucus penetrating).

**Figure S2.**
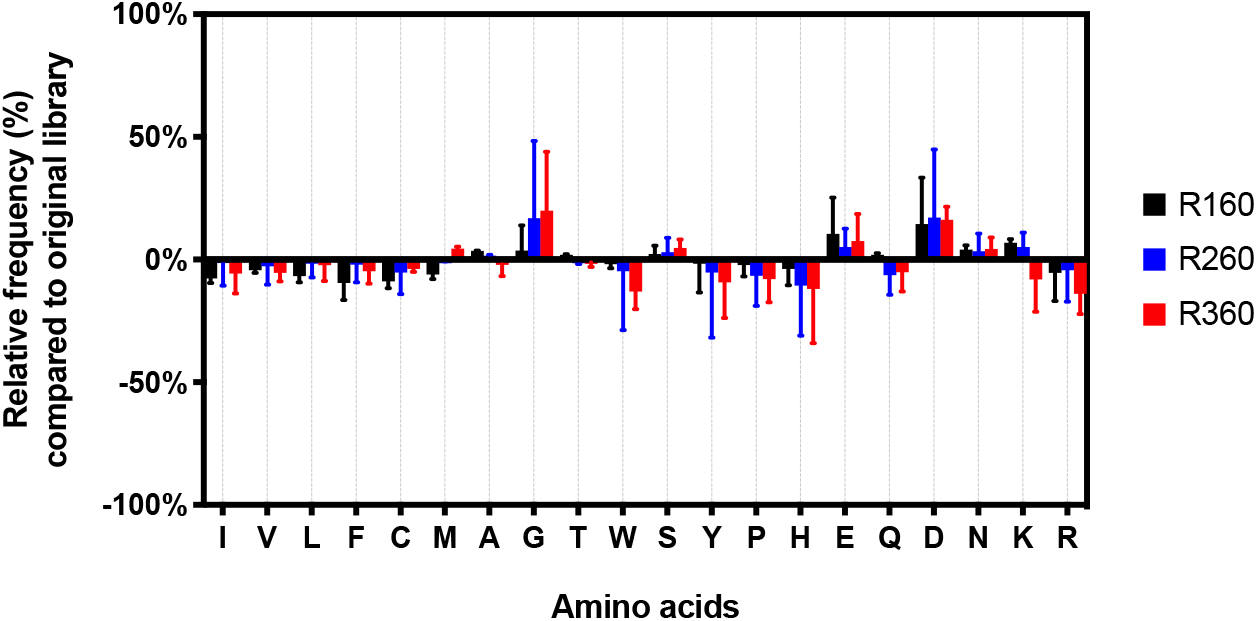
Analysis of changes in amino acid composition over rounds of selection. Overview of average amino acid composition changes (%) over rounds of all unique peptides sequences compared to the naïve library composition. Standard amino acid one-letter codes are used. Amino acids scale ranges from most hydrophobic to most hydrophilic residues, according to calculated hydropathy scores in the literature (1).

### 2. SI Tables

**Supplementary Table 1.**
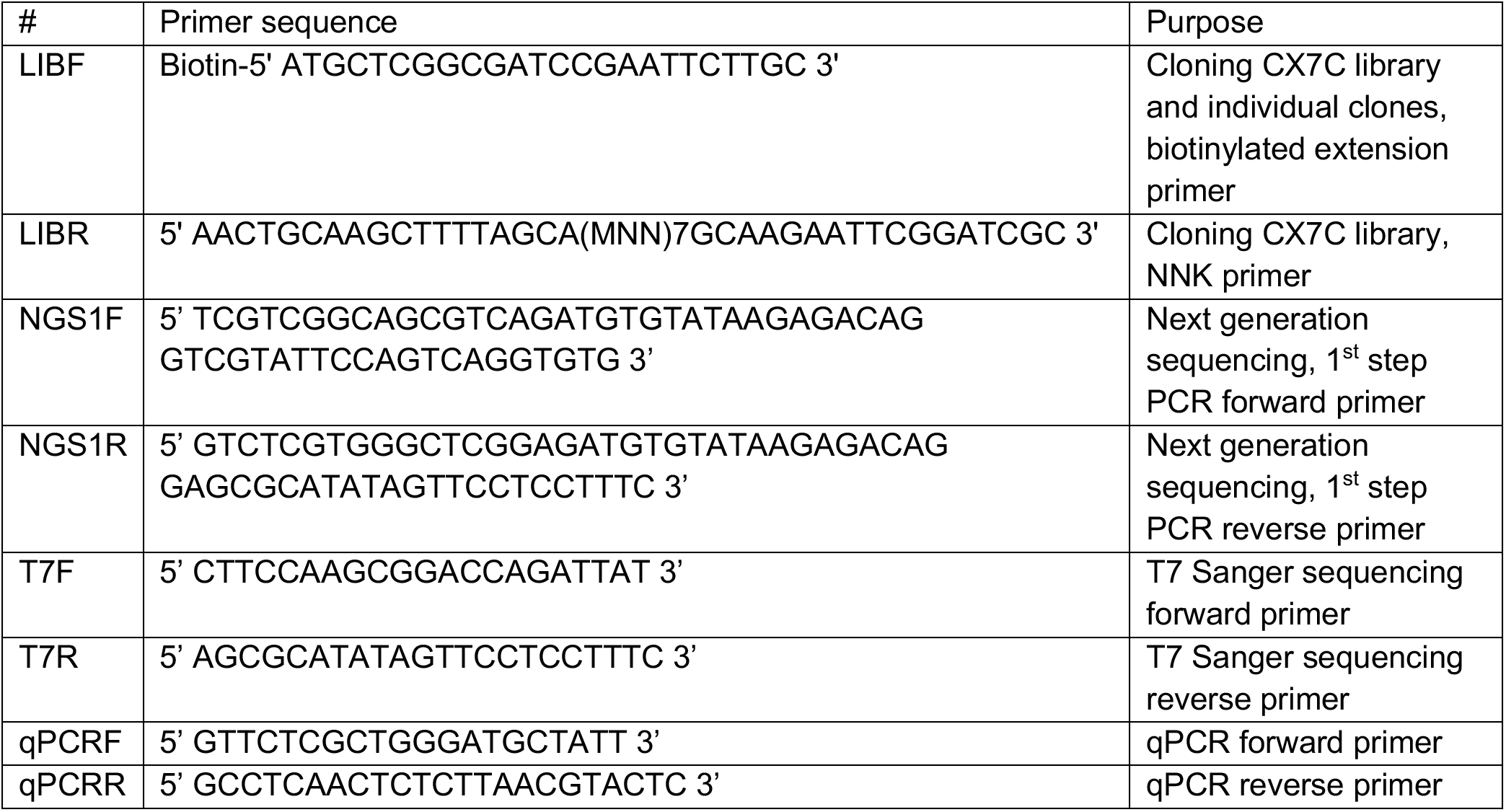
Primer Sequences

**Supplementary Table 2.** Number of sequences in each step of the processing. Provided numbers indicate the number of sequences in each particular step. *without selection, just amplification.

| Sample | Selection round | Number of total reads | Unique peptides |
| --- | --- | --- | --- |
| R115N1 | 1 | 548503 | 62233 |
| R115N2 | 1 | 631834 | 30381 |
| R130N1 | 1 | 541356 | 59001 |
| R130N2 | 1 | 850324 | 117761 |
| R160N1 | 1 | 816262 | 296972 |
| R160N2 | 1 | 766299 | 131432 |
| R215N1 | 2 | 670998 | 6581 |
| R215N2 | 2 | 638945 | 3696 |
| R230N1 | 2 | 745741 | 11298 |
| R230N2 | 2 | 631443 | 2978 |
| R260N1 | 2 | 678387 | 36364 |
| R260N2 | 2 | 613838 | 37641 |
| R315N1 | 3 | 716516 | 2862 |
| R315N2 | 3 | 677479 | 3496 |
| R330N1 | 3 | 590962 | 9167 |
| R330N2 | 3 | 444176 | 2355 |
| R360N1 | 3 | 739228 | 20328 |
| R360N2 | 3 | 502589 | 19178 |
| T7 library | 0* | 584850 | 369086 |
| T7libR1 | 1* | 290441 | 125856 |
| T7libR2 | 2* | 731956 | 268379 |
| T7libR3 | 3* | 557957 | 90786 |

**Supplementary Table 3.**
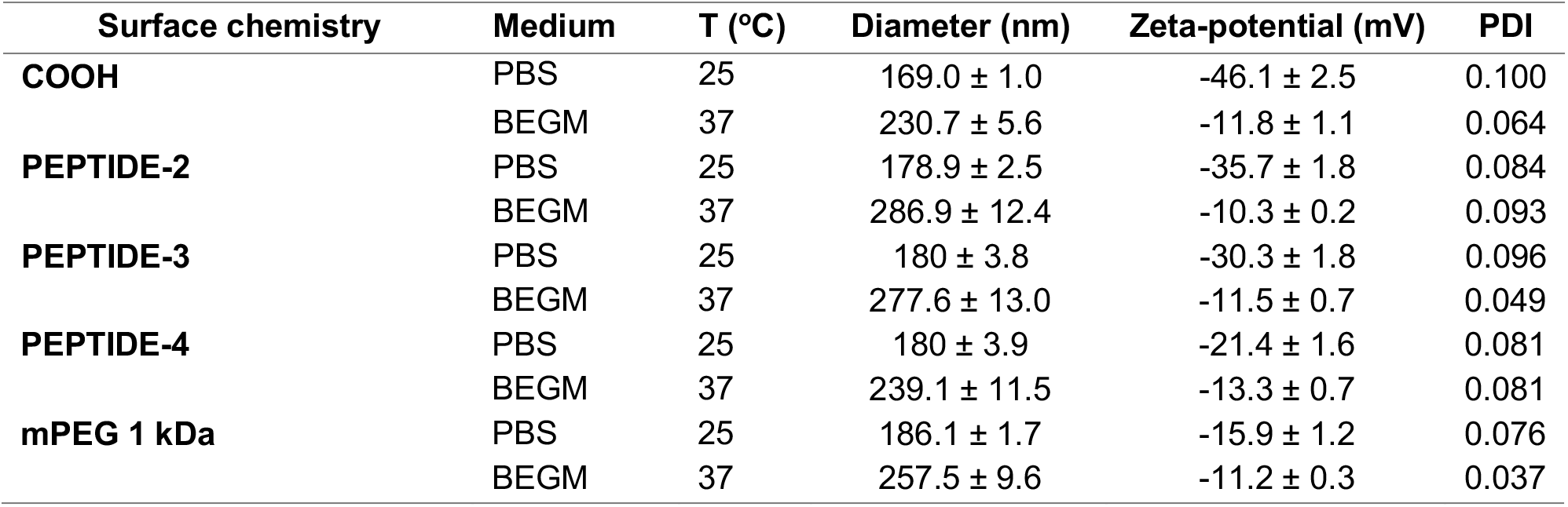
Characterization of fluorescent carboxyl-modified 100 nm nanoparticles conjugated onto their surface via EDC chemistry with peptides sequences 2-4, and mPEG 1KDa measured by DLS in 1X PBS at 25 °C and BEGM media at 37°C. Data represents mean ± SD (n = 3).

